# The Arabidopsis nucleoporins 98A/B are essential for NLP7 nuclear accumulation in response to nitrate

**DOI:** 10.1101/2025.09.22.677877

**Authors:** Zsolt Kelemen, Garry Duville, Virginie Bréhaut, Anne Krapp

**Author notes:** These authors have equally contributed (Zsolt Kelemen, Garry Duville). Anne Krapp, Institute Jean-Pierre Bourgin for Plant Sciences (IJPB), Route de Saint Cyr, 78000, Versailles, France, **Email:**.

## Abstract

The nuclear compartment is an essential evolutionary gain in eukaryotes. Stimulus-induced nuclear accumulation of transcription factors allows rapid and reversible regulation of gene expression. Nitrate-regulated nuclear accumulation of the master transcriptional regulator NLP7, a member of the RWP-RK family of transcription factors, triggers a major gene expression cascade. Whether and how this process is regulated at the level of the nuclear pore remains elusive. Here we show that the mobile nucleoporin NUP98A and its close homolog NUP98B interact with NLP7 in the nucleus and that the nitrate-triggered nuclear accumulation of NLP7 is impaired in their absence. We also demonstrated that the simultaneous loss of NLP7 and NUP98A has a strong and specific effect on the expression of nitrate-regulated genes, resulting in a drastic reduction in plant growth. Our results indicate that plant NUP98 acts on nitrate use by plants through its interaction with the top-tier NLP7 transcription factor.

**Significance Statement:** Nucleoporins (NUPs) are fundamental components of all eukaryotic cells. They are increasingly recognized not only as structural components of the nuclear pore complex but also as active regulators of gene expression and signaling pathways across eukaryotes. NUP98 proteins in mammals are multifunctional proteins with key roles beyond nucleocytoplasmic transport. The adaptation of plants to nitrogen availability is crucial for their development and yield. The master transcription factor NLP7 orchestrates early responses to nitrate supply and is regulated by a rapid nitrate-regulated nucleo-cytosolic shuttling. In this study, we demonstrate that the Arabidopsis nucleoporins 98A/B enable acclimation of Arabidopsis to N status by gating the nucleocytoplasmic trafficking of the nitrate sensor NLP7 transcription factor, enlarging their functional network.

**Graphical abstract:** 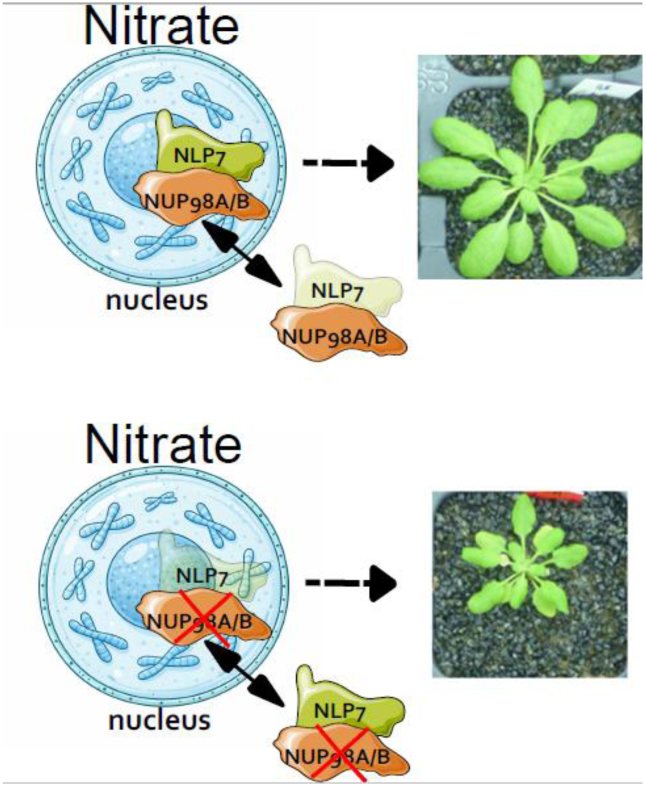

## Introduction

Nucleoporins (NUPs) are fundamental components of all eukaryotic cells. They are constituents of the nuclear pore complexes (NPCs), which are extremely large protein complexes embedded in the nuclear envelope and consist of over 1,000 copies of ∼30 individual NUPs. The central channel of the NPC functions as a permeability barrier between cyto- and nucleoplasm. The FG-repeat-containing NUPs within the central channel constitute a permeability barrier that allows the passive diffusion of small molecules, while limiting the passage of macromolecules larger than ∼40 kDa and enabling nucleocytoplasmic exchange. It thus plays the role of a regulator of intracellular molecule trafficking, constituting an important gatekeeper for macromolecular transports between the nucleus and cytoplasm (Li et al., 2016).

In addition to their main function in nucleocytoplasmic transport, NPCs and their components have been shown to play important roles in many biological processes, such as kinetochore and spindle assembly, cell cycle control, regulation of gene expression, chromatin organization, DNA repair, and DNA replication (Capelson and Hetzer 2009; Capelson et al., 2010; Strambio-De-Castillia et al., 2010; Wozniak et al., 2010; Van de Vosse et al., 2011; Bukata et al., 2013). In Arabidopsis, NUPs are also involved in multiple biological processes, including flowering and immune responses (Gu et al., 2016; Cheng et al., 2020).

Changing environmental conditions necessitate rapid adaptation of cytoplasmic and nuclear protein content, and a number of key transcription factors are regulated by a rapid nuclear accumulation in response to a signal (Cartwright and Helin, 2000). These TFs must be transported from the cytoplasm to the nucleus to reach their targets and induce or repress their transcription, and exported from the nucleus to the cytoplasm.

Nitrogen (N) is a crucial macronutrient, and N availability in the soil can be a limiting factor for plant growth and development (Marschner et al., 1995). Thus, plants evolved mechanisms to sense and promptly adapt to fluctuating nitrate availability. Importantly, nitrate is not only a nutrient but also a signaling molecule (Crawford, 1995; Scheible et al., 1997). The rapid transcriptional response to nitrate supply triggers expression changes of thousands of genes.

Members of the Nin Like-Protein (NLP) family are top tier TFs of the nitrate signaling pathway through inducing a cascade of transcriptional regulation (reviewed in Mu and Luo, 2019). The Arabidopsis NLP7 TF is regulated by a very rapid nuclear accumulation within seconds after exposure to nitrate. This mechanism involves the phosphorylation of an evolutionarily conserved Serine by three CALCIUM-DEPENDENT PROTEIN KINASEs (CPK10, CPK30, CPK32, Liu et al., 2017). In addition, NLP7 activity is directly regulated by nitrate binding to its N-terminal region (Liu et al., 2022). The Arabidopsis genome encodes nine NLPs that bind in yeast the nitrate-responsive cis-element (NRE), a conserved DNA sequence motif found in the proximal regions of promoters of nitrate-responsive genes and shown to be necessary and sufficient for their nitrate-induced gene expression (Konishi and Yanagisawa, 2011, 2013). NLP7 was shown as a major TF governing the early nitrate response (Marchive et al., 2013; Konishi and Yanagisawa, 2019; Alvarez et al., 2020). NLP6, the closest homolog of NLP7, exhibits a partially redundant function to that of NLP7. However, although the loss of NLP7 is sufficient to cause a growth reduction under ample nitrate supply and impaired induction of nitrate-responsive gene, the additional loss of function of NLP6 leads to a stronger phenotype than that of the single mutant *nlp7-1* (Guan et al., 2017, Cheng et al., 2023). Recent studies demonstrated that NLP2 orchestrates both N and carbon assimilation in response to external nitrate availability (Konishi et al., 2021; Durand et al., 2023). Other NLPs also contribute to the regulation of nitrate-inducible gene expression and nitrate-dependent promotion of vegetative growth in Arabidopsis (Konishi et al., 2021; Liu et al., 2022). The functionality of NLPs is further enriched by their ability to form homo-, heterodimer, or to interact with other proteins via their PB1 domain (Guan et al., 2017; Konishi et al., 2021).

By exploring NLP7 interacting proteins, we discovered that nucleoporins NUP98A and NUP98B interact directly with NLP7. Here, our goal was to elucidate how NUP98A/B bridges nitrate signaling to the NPC function. We first confirmed the interaction of NUP98A/B with NLP7 in planta. Then, applying genetics, we showed that the double mutant *nup98anup98b* has a higher sensitivity to nitrate limitation and that the loss of function of NUP98A in the *nlp7* mutant background leads to a strongly impaired early transcriptional nitrate response and impaired growth on nitrate fertilization. Finally, the significant reduction of GFP-NLP7 nuclear accumulation in response to nitrate in the genetic background of *nup98anup98b* strongly underscores the critical role of NUP98A/B proteins in regulating nitrate-dependent nuclear accumulation of NLP7.

## Results

### NLP7 protein interacts with the nucleoporin NUP98A/B

Using a Y2H approach (performed by Hybrigenics), we found an interaction between the full-length NLP7 protein and NUP98A. We then confirmed this interaction by pairwise Y2H assay (Figure 1A) and in planta by co-expressing ^C^YFP-NLP7 and ^N^YFP-NUP98A in *Nicotiana benthamiana*. The YFP signal indicating an interaction between the co-expressed proteins was present in the nucleus as well as in the cytoplasm (Figure 1B-G).

**Figure 1.**
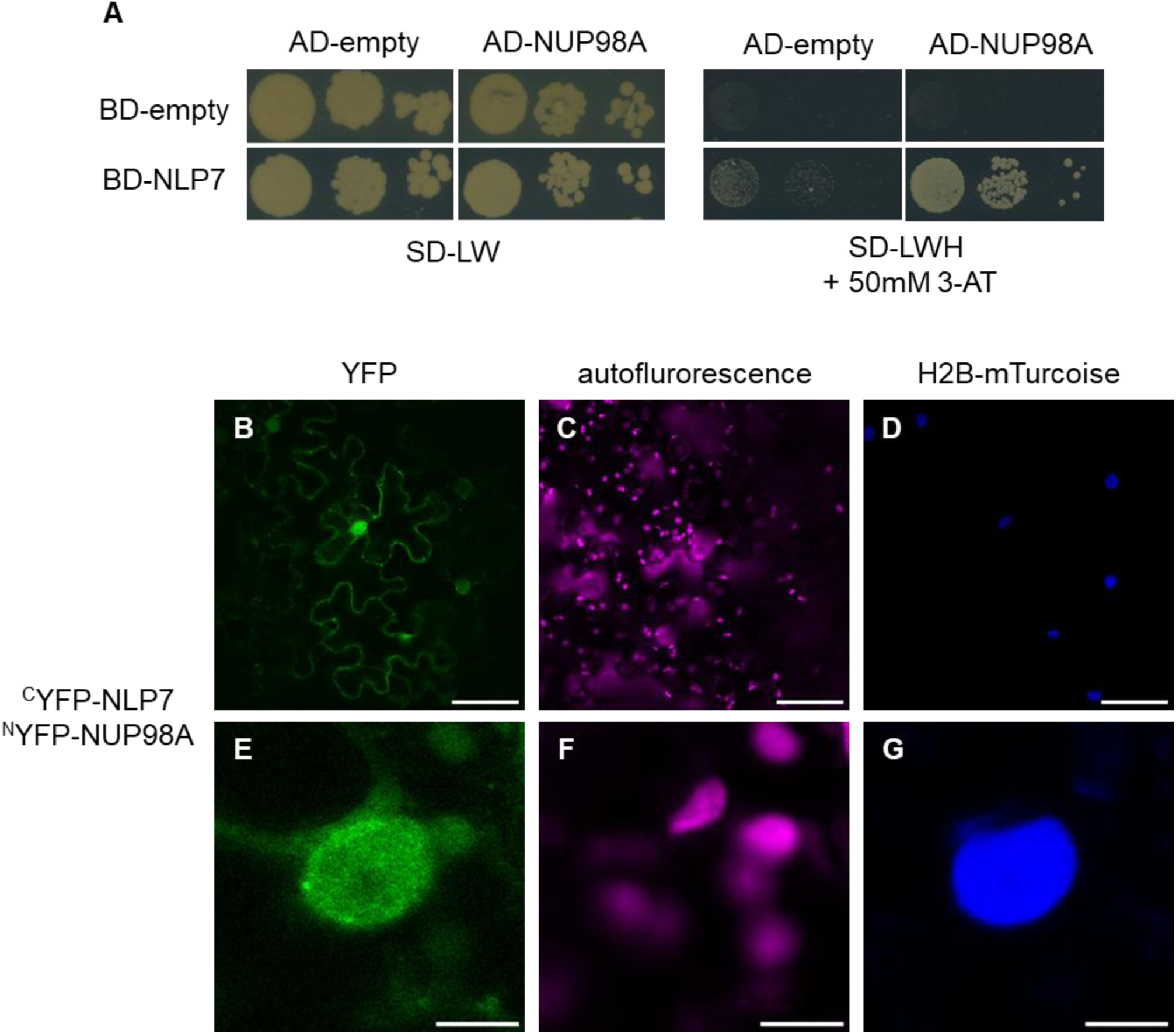
NLP7 interacts with NUP98A. (**A**) Yeast two-hybrid assay using NLP7 fused with the binding domain (BD) and NUP98A fused with the DNA activation domain (AD). Cells were grown on selective and non-selective plates. (**B-G**) NLP7 interacts with NUP98A in Bimolecular Fluorescence Complementation (BiFC) assays in *N. benthamiana*. (**B, E**) YFP fluorescence (green), (**C, F**) chlorophyll autofluorescence (magenta), (**D, G**) H2B nuclear marker - mTurcoise fluorescence (blue) Scale bar = (**B-D**) 50 µm, (**E-F**) 5 µm

We then tested whether the closely related nucleoporin NUP98B also interacts with NLP7 after transient expression in *N. benthamiana*, using two other nucleoporins, NUP96 and CPR5, as negative controls (Figure 2). NLP7 showed interaction with NUP98B with a localization of the interaction similar to NUP98A (Figure 2D-F), whereas the nucleoporins NUP96 and CPR5 showed no interaction with NLP7 (Figure 2G-L). Taken together, these results demonstrate that the interaction between NLP7 and NUP98A/B is specific to these two NUPs and is not a general interaction with proteins of the NPCs.

**Figure 2.**
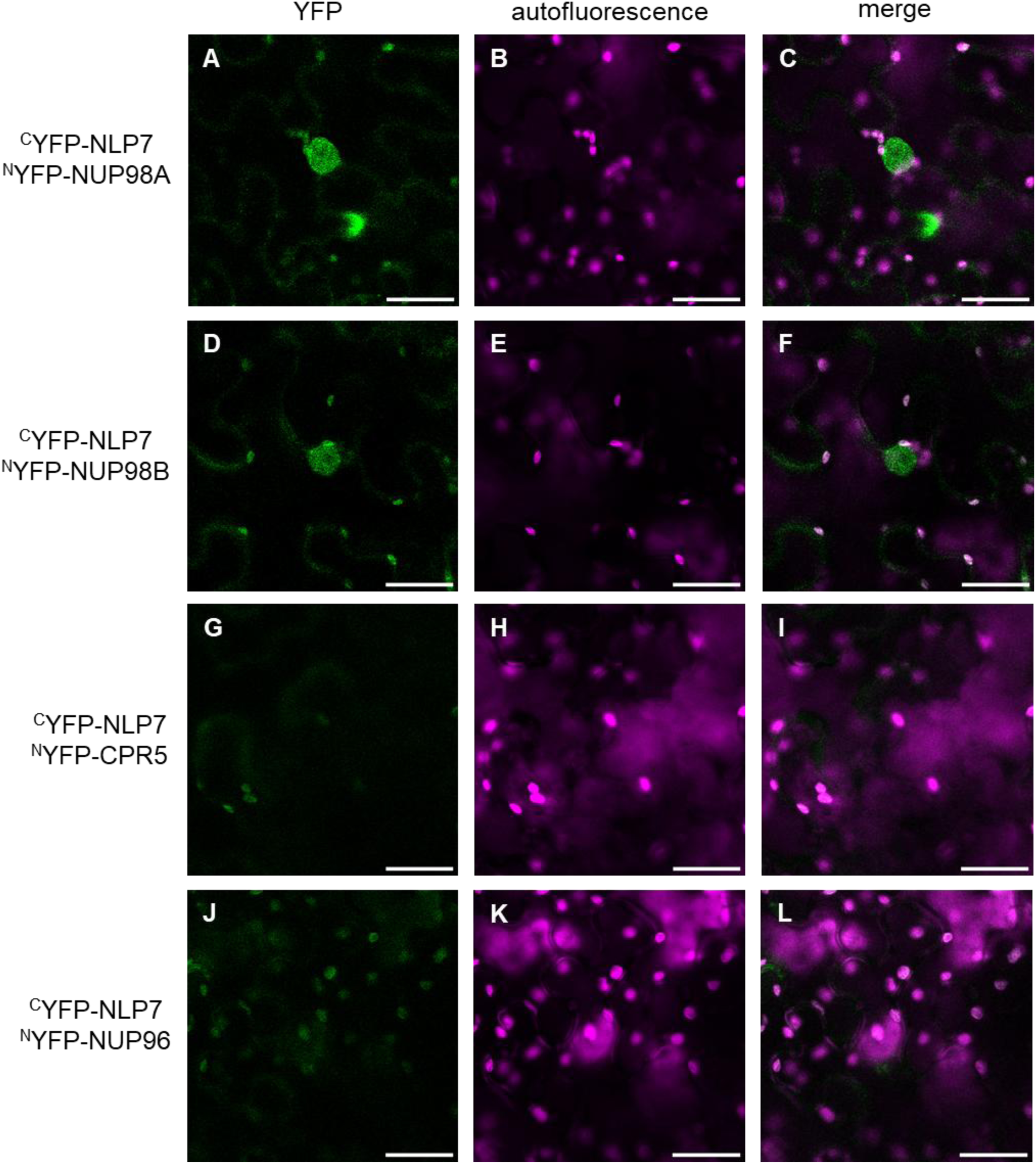
NLP7 interacts with NUP98A and NUP98B. **(A-L)** NLP7 interacts with NUP98A and NUP98B in BiFC assays in *N. benthamiana.* (**A-C**) NLP7-NUP98A interaction, (**D-F**) NLP7-NUP98B interaction, **(G-I)** NLP7-CPR5 interaction (negative control), **(J-L)** NLP7-NUP96 interaction (negative control). (**A, D, G, J**) YFP fluorescence (green), (**B, E, H, K**) chlorophyll autofluorescence (magenta) (**C, F, I, L**) YFP fluorescence (green) was merged with chlorophyll autofluorescence (magenta). Scale bar = 25 µm.

### PB1 and GAF domains of NLP7 are not essential for NLP7-NUP98A interaction

NLPs contain 2 predicted protein-protein interaction domains. The C-terminal PB1 domain, which has been shown to be involved in the formation of homo- and heterodimers of NLP proteins (Konishi and Yanagisawa, 2019) and with proteins that do not contain PB1 domains (Guan et al., 2017). The second, the N-terminal GAF-like domain, is reported as a protein-protein interacting domain (Ho et al., 2000; Wu et al., 2022). To identify the NLP7 protein domains involved in the interaction with NUP98A, we produced NLP7 constructs without either the GAF-like domain or the PB1 domain. NLP7 without either GAF or PB1 domain interacted with NUP98A in the BIFC assays, producing positive signal both in the nucleus and in the cytoplasm (Supplemental Figure S1). Thus, the NLP7 PB1 and GAF domains might contribute to the interaction between NLP7 and NUP98A/B but are not essential for it.

### The intracellular localization of the NUP98A-NLP7 interaction is modified by the N status of the plants

NLP7 accumulates in the nucleus in the presence of nitrate and is localized in the cytosol in N-starved seedlings (Marchive et al., 2013). Because NUP98A/B are key components of the NPC, we then ask whether their association with NLP7 might depend on the N status. We studied the interaction between NLP7 and NUP98A in *N. benthamiana* plants that have been grown under ample nitrate supply (10 mM) and in plants that were treated with a 5-day N depletion. In addition, leaf discs from N-depleted plants were resupplied with nitrate 10 min before observation (Figure 3). The fluorescent signal indicating the interaction between NLP7 and NUP98A was present both in the cytoplasm and in the nucleus in all three conditions, however, in the case of the starved leaves, the nuclear signal was less defined (Figure 3 D-F). Quantification of the fluorescence ratio between nucleus and cytoplasm confirmed these observations (Figure 3 J). Our results suggest that NUP98A/B interact with NLP7s when both proteins are localized in the same cellular compartment with enhanced interaction in the nucleus compared to the cytosol in the presence of nitrate.

**Figure 3.**
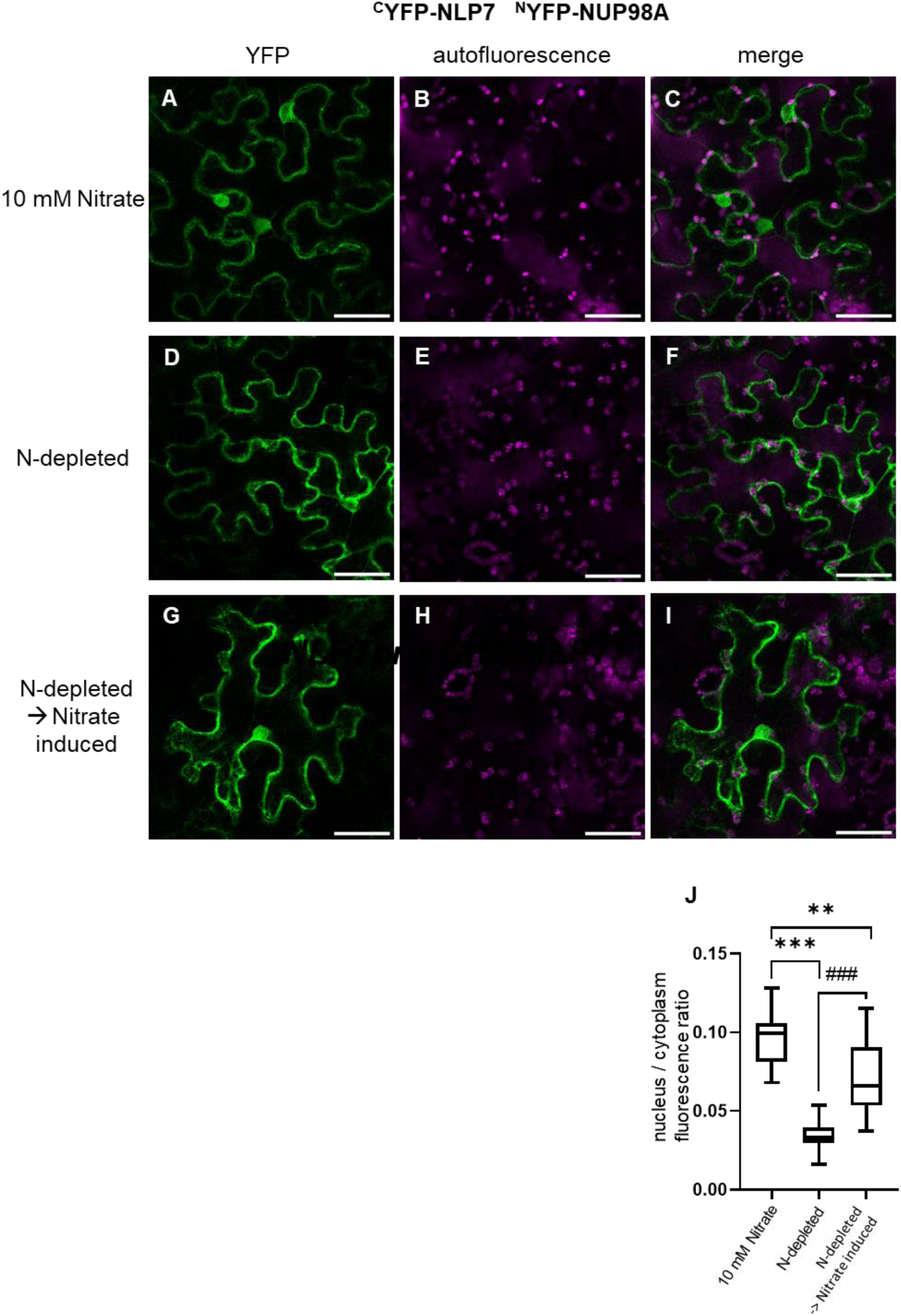
NLP7 interacts with NUP98A in the nucleus in the presence of nitrate. **(A-1)** BiFC assays between NLP7 and NUP98A in *N. benthaminia.* **(A-C)** Plants were grown under ample (10 mM) nitrate nutrition **(D-F)** Plants have been N-depleted for 5-days **(G-1)** leaf disc from transitory transformed N-depleted plants were treated for 10 min with nitrate. (A, D, G) YFP fluorescence (green), (B, E, H) chlorophyll autofluorescence (magenta) (C, F, I) YFP fluorescence (green) was merged with chlorophyll autofluorescence (magenta). Scale bar = 50 µm. **(J)** Nuclear-to-cytoplasmic fluorescence intensity ratios in plant cells across different samples. Ratios were calculated from measurements in 13 - 15 images per genotype and treatment. Data are presented as boxplots. Statistical significance was assessed using the Kruskal-Wallis test followed by Dunn’s post hoc test for pairwise comparisons. P-values: 4.47e-02 (**), 2.7e-07 (***), 1.08e-03 (###).

### Limiting nitrate supply amplifies the early senescence of nup98anup98b double mutants

The protein interaction between NLP7 and NUP98A/B led us to evaluate the role of NUP98A/B in the response of plants to nitrate availability. We analyzed the growth of NUP98A/B loss-of-function mutants (*nup98a-1*, *nup98a-2*, *nup98b-1*, *nup98b-2* and *nup98a-1nup98b-1*) under contrasting nitrate supply that had revealed nitrate-dependent growth phenotypes of *nlp7* and *nlp2* mutants (Durand et al., 2023). We grew the plants in sand culture under short-day conditions (8 h light/16 h dark) with either 0.5 mM nitrate or 5 mM nitrate supply for 39 days and determined rosette and root biomass (Figure 4A - D). Under both N supply conditions, the rosette and root biomass of *nup98a* and *nup98b* single mutants were similar to that of the wild type (WT) Col-0. Under non-limiting N supply, WT plants displayed the expected (2.5-fold) increase in rosette fresh weight compared to WT plants grown under 0.5 mM N supply (Figure 4A and B). Rosette biomass of the double mutant *nup98a-1nup98b-1* was strongly decreased in both N regimes (by 72 and 84 % under 0.5 and 5mM nitrate supply, respectively). Root fresh weight of *nup98a* and *nup98b* mutants was also similar to WT, with a trend towards slightly lower root biomass compared to WT under limiting N supply (Figure 4 C and D). Root biomass was also decreased in the case of the *nup98a-1nup98b-1* double mutant, and the rosette/root ratio was modified for the double mutant compared to WT (Figure 4 E and F). Interestingly, it decreased slightly by 30% under non-limiting N supply, but increased 2-fold under limiting N supply. The shoot/root ratio is an indicator of N limitation, suggesting that the double mutant experienced less N limitation than the WT and the single mutants under limiting N supply. Remarkably, the early senescence phenotype of *nup98a-1nup98b-1* double mutant that was described before for this double mutant (Xiao et al., 2020) and that was visible by the loss of chlorophyll, particularly in older leaves, had appeared much earlier in plants grown under nitrate-limiting conditions than under ample nitrate supply (at 28, 35 and 38 days, respectively, Figure 4, G and H). Under these conditions no loss of chlorophyll was observed for WT under both nitrate regimes. Taken together, these observations suggest that NUP98A/B are necessary to prevent early N limitation-induced senescence.

**Figure 4.**
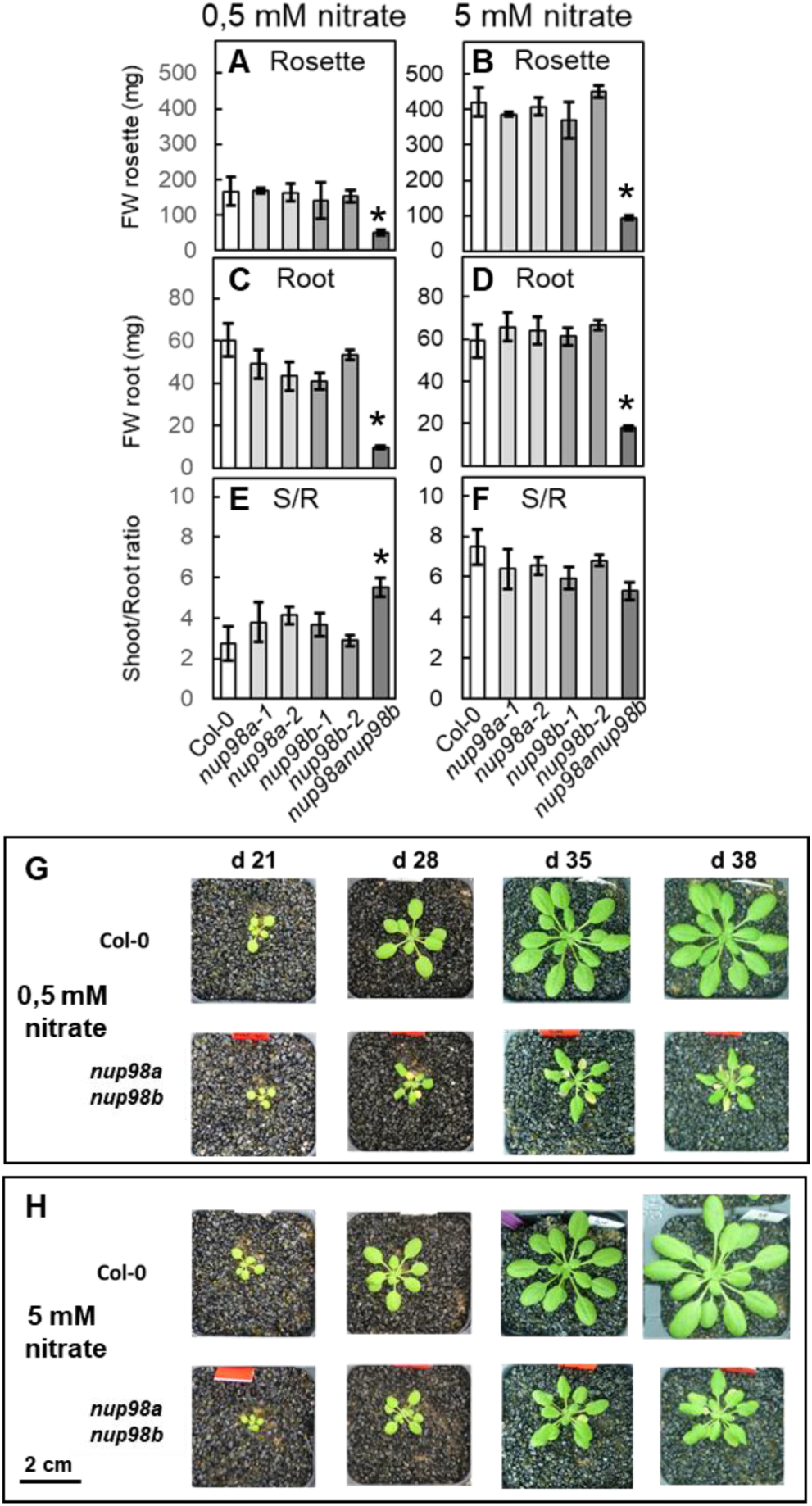
Biomass and Shoot/Root ratio of *nup98a/b single and double* mutants grown under limiting and non-limiting nitrate supply. Plants were grown on sand under short days with a light intensity of 150 µE for 39 d (**A, B**) Rosette fresh weight of plants grown under limiting (0.5 mM, **A**) and non-limiting (5 mM, **B**) nitrate supply. (**C, D**) Root fresh weight of plants grown under limiting (0.5 mM, **C**) and non-limiting (5 mM, **D**) nitrate supply. (**E, F**) Shoot/root ratio of plants grown under limiting (0.5 mM, **E**) and non-limiting (5 mM, **F**) nitrate supply. Data were obtained from six plants from two independent experiments. Significant differences between genotypes for each nitrate supply condition were determined by Student t-Test (* *P*-value ≤ 0.05). **(G, H)** Top view of rosettes of Col-0 and *nup98anup98b-*double mutants grown under limiting (0.5 mM, **G**) and non-limiting (5 mM, **H**) nitrate supply at different times of the culture. Scale bar = 2 cm.

### Double and triple mutant phenotypes support an interplay between NLP7 and NUP98A/B

To study the genetic interaction of NUP98A/B and NLP7 and to further unravel the roles played by NUP98A/B in the NLP7-dependent nitrate responses, we generated the *nup98a-1nlp7-1* and *nup98b-1nlp7-1* double mutants and the triple mutant *nup98a-1nup98b-1nlp7-1*.

We first studied the growth of single and higher order mutants in comparison to the WT in sand culture under low and ample nitrate supply in short day conditions (Figure 5 A-C). Under limiting N supply, the single mutants *nup98a*, *nup98b* and *nlp7-1* attained a rosette biomass comparable to that of the WT, as expected. The rosette FW of the double mutants *nlp7-1nup98a* and *nlp7-1nup98b* decreased by 40% and 30% respectively, compared to *nlp7-1*. The double mutant *nup98anup98b* had a rosette biomass of 37 mg, corresponding to 22% of WT and the triple mutant *nup98a-1nup98b-1nlp7-1* barely attained an FW of 1 mg (Figure 5A). Under non-limiting nitrate supply, whereas the *nup98a* and *nup98b* showed no difference to WT, the biomass of the *nlp7-1* mutant was decreased 2-fold compared to WT, as reported previously and explained by a constitutive N-limitation phenotype under ample nitrate supply (Castaings et al., 2009). In these conditions, rosette biomass of the double mutants *nlp7-1nup98a* and *nlp7-1nup98b* was decreased by 50% and 35%, respectively, compared to *nlp7-1*, thus slightly stronger than under limiting nitrate supply. The triple mutant reached a biomass of 5.5 mg compared to 75 mg for the double mutant *nup98anup98b*. Together, our data suggest that both NLP7 and NUP98A/B determine plant growth in response to N supply. This can be due either to their participation in a shared regulatory pathway or in two different pathways affecting growth in response to the N status of the plant.

**Figure 5.**
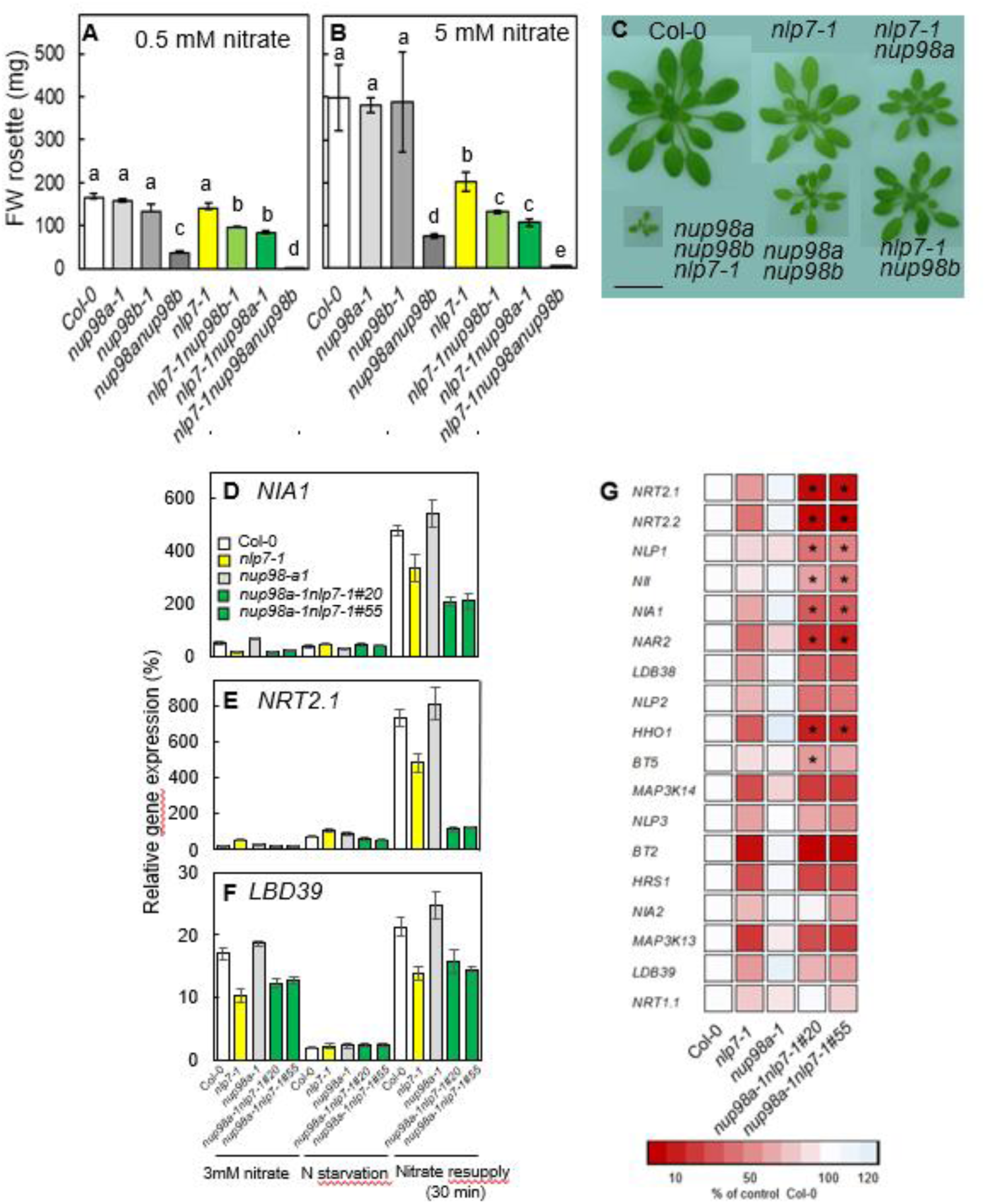
Growth and gene expression in response to nitrate are strongly impacted by the simultaneous loss of function of Nucleoporin 98A/B and NLP7. **(A-C**) Plants were grown on sand under short days with a light intensity of 150 µE for 39 days (**A, B**) Rosette fresh weight of plants grown under limiting (0.5 mM, **A**) and non-limiting (5 mM, **B**) nitrate supply. **(C)** Top view of rosettes of plants grown under non-limiting (5 mM) nitrate supply, Scale bar = 2 cm. **(D-F)** Gene expression analysis. Relative transcript levels of selected genes before and after nitrate addition to N-starved 14-day-old seedlings. Expression was measured by RT-qPCR and normalized as a percentage of the expression of a constitutive synthetic gene composed of *TIP41* (At4g34270) and *ACTIN2* (*ACT2*, At3g18780). **(G)** Heatmap representation of the fold-change in steady state transcript level after 30 min of nitrate resupply in comparison to Col-0. Red, lower abundance; blue, higher abundance (* *P*-value ≤ 0.05 in comparison to *nlp7-1*).

### Loss of function of both NLP7 and NUP98 amplified the altered early response to nitrate in nlp7-1 mutants

Given the conserved role of nucleoporins and transcription factors in many eukaryotes, the interaction between NLP7 and NUP98A/B may reflect a broader principle whereby nuclear transport components cooperate with transcriptional regulators to fine-tune N responses. We compared steady state levels of mRNA in seedlings either grown for 11 days on 3 mM nitrate medium, or deprived of nitrogen for 3 days after 11 days in medium containing 3 mM nitrate, or resupplied for 30 min by 3 mM nitrate after the 3-day deprivation period. We measured by RT-qPCR the relative transcript levels of 18 NLP7-dependent nitrate-regulated genes in *nlp7-1nup98a* double mutants compared to the single mutants and the WT (Figure 5 D-G). The main differences between *nlp7-1* and the double mutant *nlp7-1nup98a* have been observed after nitrate resupply (Figure 5 G). For all of the studied genes, the nitrate-triggered increase in steady-state transcript level was diminished for *nlp7-1* as expected. For the double mutant *nlp7-1nup98a*, this reduced increase was further diminished in the case of *NITRATE TRANSPORTER 2.1* (*NRT2.1*), *NRT2.2*, *NLP1*, *NITRITE REDUCTASE* (*NII*), *NITRATE REDUCTASE1* (*NIA1*), *NAR2*, and *HRS1 HOMOLOGOUS 1* (*HHO1*). A similar tendency was observed for *LOB BINDING DOMAIN 38* (*LBD38*), *NLP2* and *BTB/POZ AND TAZ DOMAIN-CONTAINING PROTEIN 5* (*BT5*), whereas no difference in the expression level after 30 min of nitrate resupply for the double mutant *nlp7-1nup98a* compared to *nlp7-1* was found for *MAP KINASE KINASE KINASE 14* (*MAP3K14*), *MAP3K13*, *NLP3*, *BT2*, *HYPERSENSITIVITY TO LOW PI-ELICITED PRIMARY ROOT SHORTENING 1* (*HRS1*), *LBD39*, *NIA2* and *NRT1.1*. Taken together, for several nitrate-regulated genes, the combination of *nlp7* and *nup98* mutations resulted in an additive decrease in nitrate induction, likely due to the consequences of impaired nuclear accumulation of other NLPs or other transcription factors necessary for complete early response to nitrate.

### NUP98A and NUP98B are required for the nitrate-triggered nuclear accumulation of NLP7

To decipher the mechanism underlying the various developmental and molecular phenotypes of the higher-order mutants, we studied the nuclear accumulation of GFP-NLP7 in response to nitrate resupply to N-depleted seedlings in different mutant backgrounds. We hypothesized that the interaction of NUP98A and NUP98B with NLP7 is required for the nitrate-triggered nuclear accumulation and NLP7-dependent nitrate-regulated gene expression and growth.

We introduced the *GFP-NLP7* translation fusion driven by the *NLP7* promoter into these NUP98 loss of function mutants by crossing an *NLP7pro:GFP-NLP7* (*nlp7-1*) transgenic line (Castaings et al., 2009) with the single mutant *nup98a-1* and the double mutant *nup98a-1nup98b-1*. In N-depleted 10-day-old seedlings, GFP-NLP7 was localized in the cytosol of all genotypes (Figure 6 A, C, E). At 10 minutes after nitrate resupply, in the presence of NUP98A and NUP98B GFP-NLP7 accumulated rapidly in the nucleus (Figure 6 B). The loss of function of NUP98A was accompanied by a decrease in the accumulation in the nucleus after nitrate resupply (Figure 6 D). In the absence of both NUP98A and NUP98B, the accumulation of GFP-NLP7 in the nucleus in response to nitrate resupply was almost abolished (Figure 6 F). Quantification of the fluorescence ratio between nucleus and cytoplasm confirmed these observations (Figure 6 G). Thus, our results indicate a specific role of NUP98A/B for the nitrate-triggered nuclear accumulation of NLP7.

**Figure 6.**
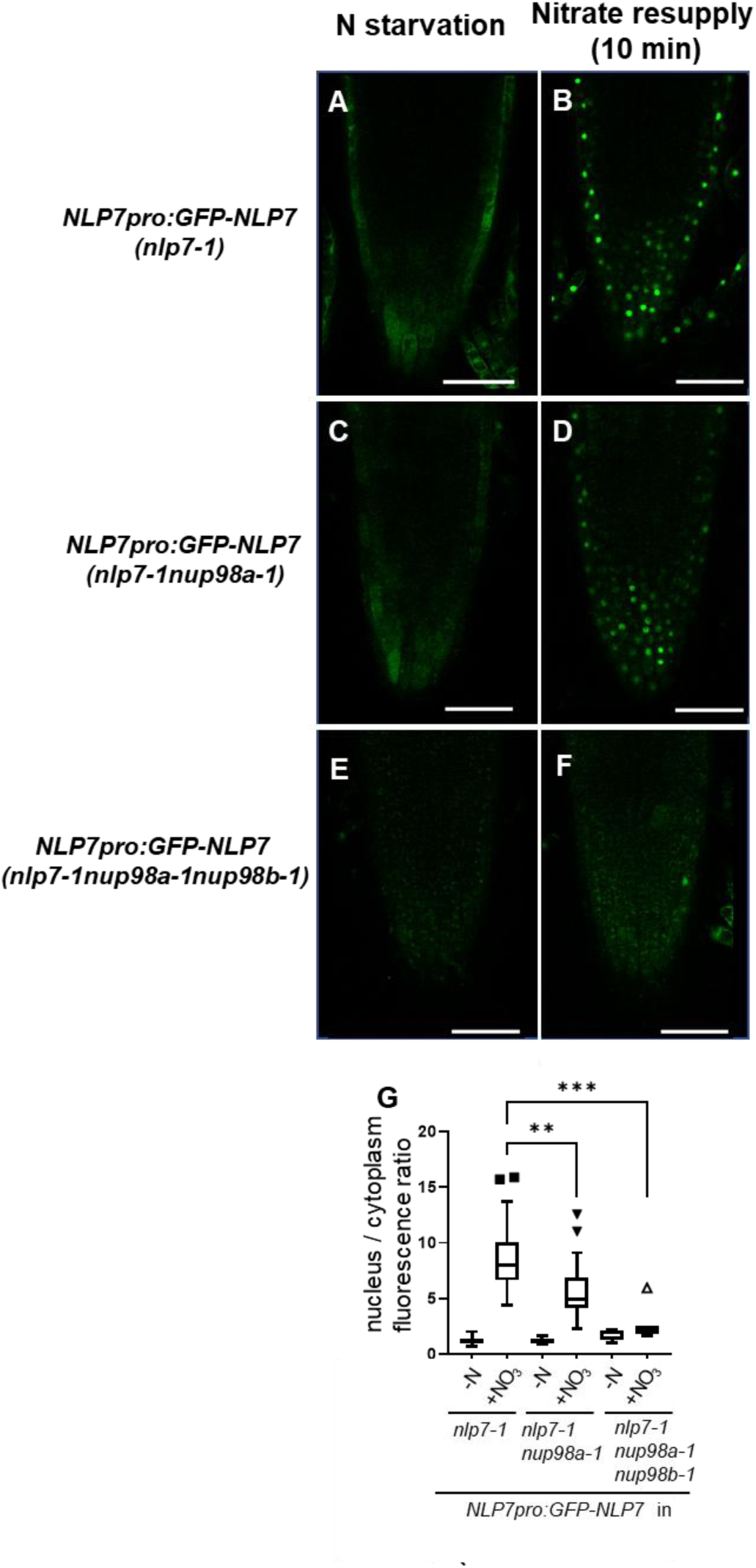
NUP98A *and* NUP98B loss of function abolished the nitrate-mediated nuclear accumulation of NLP7. Analysis of the nitrate-triggered nuclear accumulation of GFP-NLP7. **(A-H)** 10-days-old seedlings were N starved for 2 days before observation by confocal microscopy. Acquisitions were performed before nitration addition **(A, C, E),** or 10 min after nitrate addition **(B, D, F).** Representative images out of 3 independent experiments are shown. Scale bar= 50 µm. **(G)** Nuclear-to-cytoplasmic fluorescence intensity ratios in plant cells across different samples. Ratios were calculated from measurements in 7 - 15 images per genotype and treatment. Data are presented as boxplots. Statistical significance was assessed using the Kruskal-Wallis test followed by Dunn’s post hoc test for pairwise comparisons. P-values: 1.49e-3 (**), 6.5e-6 (***). Black squares (▪) and black downward triangles (▼): regular outliers; white upward triangle (△): extreme outlier.

## Discussion

Nucleoporins are increasingly recognized not only as structural components of the nuclear pore complex (NPC) but also as active regulators of gene expression and signaling pathways across eukaryotes. NUP98 proteins in mammals are multifunctional components of the nuclear pore complex with key roles beyond nucleocytoplasmic transport. Increasing evidence shows that NUP98 and other nucleoporins are frequently associated with active chromatin regions, particularly enhancer elements, where they contribute to the regulation of gene expression and chromatin organization. NUP98 can act as a scaffold at these regulatory sites, promoting the maintenance of transcriptionally active chromatin by stabilizing histone modifications and facilitating epigenetic memory. Several models have been proposed to explain this chromatin-binding function of NPCs: (i) NPCs may help maintain the active state of genes through direct interaction with histone-modifying enzymes; (ii) the targeting of active genes and enhancers to the NPC could be functionally linked to transport-related mechanisms, such as the nuclear shuttling of transcription factors; and (iii) the intrinsically disordered FG-repeat domains of NUP98 may enable phase separation at NPCs, creating specialized transcriptional compartments that enhance gene activation at enhancers and super-enhancers (for review, see Pascual-Garcia & Capelson (2019). In this study, we uncover a novel role for the plant nucleoporins NUP98A and NUP98B in controlling the nuclear accumulation of the transcription factor NLP7, a central regulator of nitrogen signaling. While NUP98 proteins are generally thought to maintain NPC integrity and mediate nucleo-cytosolic trafficking (Tamura et al., 2010; Raveh et al., 2016), studies in yeast, animals, and plants have highlighted their mobility and functional versatility beyond nuclear transport (Capitanio et al., 2017; Gallemí et al., 2016). Our work now extends this paradigm to nitrogen signaling in plants.

Here, we report a novel role for the nucleoporins NUP98A and NUP98B in regulating the nuclear accumulation of the TF NLP7. Based on their sequence homology with other species, plant NUP98A/B are considered to be required for maintaining the NPC structure and facilitating macromolecule nucleo-cytosolic trafficking (Tamura et al., 2010; Raveh et al., 2016). Although they have an important structural role in the NPC, NUP98s in yeast, animals and plants are mobile proteins that can be localized in different cellular compartments (Capitanio et al., 2017; Gallemí et al., 2016). This has various implications for their role in other biological processes. We validated the interaction between NUP98A and NLP7 using in planta BiFC assays, confirming the results obtained through yeast two-hybrid screening. Interaction was detected both in the cytosol and the nucleus. We cannot exclude that the strong transient expression of the splitYFP proteins driven by the *CaMV 35S* promoter leads to a protein localization different from the native conditions. However, the localization of the interacting complex corresponded to the localization of NLP7 tagged with GFP under the particular condition, as in N-depletion condition, the interaction in the nucleus was less pronounced, as expected from the NLP7 localization change under this condition (Marchive et al., 2013).

Here, we showed that neither the NLP7 GAF domain nor its PB1 domain is essential for the interaction with NUP98A/B, however, it is possible that both domains act redundantly to ensure this interaction. Found in a variety of proteins, including TFs, the GAF domain is important for protein-protein interactions and can bind a variety of ligands, including cGMP, cAMP, and small molecules such as oxygen and nitrate (Ho et al., 2000; Moglich et al., 2009; Niemann et al., 2014; Su and Lagarias, 2007; Grefen et al., 2008). It has been shown that the NLP7 PB1 domain is important for its interaction with other NLPs, but also with the transcription factor teosinte branched1/cycloidea/proliferating cell factor20 (TCP20), and SUCROSE NON-FERMENTING 1-RELATED KINASE 1 (SnRK1) (Konishi and Yanagisawa, 2019; Guan et al., 2017; Wang et al., 2022). The NLP7 protein possesses short intrinsic disordered regions (IDRs), which leads to the hypothesis that these IDRs interact with the FG motifs of NUP98. However, the nature of the underlying molecular interactions between FG repeats that might lead to protein condensates (Ahn et al., 2025) is largely unknown. Interestingly, it has been shown recently that protein condensation is involved in the specificity of transport through the NPC (Wang et al., 2023).

Despite the high probability that the positive BiFC between NLP7 and NUP98A/B is due to a direct interaction between these proteins, we cannot exclude that a third partner is necessary for the interaction. For example, NirA, the specific transcription factor of the nitrate assimilation pathway of Aspergillus nidulans, interacts with the nuclear export factor KapK, which bridges an interaction with a nucleoporin-like protein. Interestingly, NirA, which has no structural homology with NLP7, also accumulates in the nucleus upon induction by nitrate via a nuclear retention mechanism (Bernreiter et al., 2007).

To further assess the biochemical mechanisms, further cell biological experiments are needed in *Arabidopsis thaliana* transgenic plants using native promoters to examine whether NUP98A/B and NLP7 co-localize and translocate together into the nucleus, particularly under nitrate treatment conditions. In addition, biochemical validation of the NLP7–NUP98 interaction under native expression levels and mapping of the interacting domains of NUP98A/B will be important to determine whether this regulation relies on direct binding to FG-repeat regions or involves additional transport factors. However, our genetic data demonstrate clearly that NUP98A/B are essentiel for nitrate-triggered nuclear accumulation of NLP7. Future work will be necessary to reveal the precise molecular mechanism and distinguish whether NUP98A/B primarily facilitate nuclear import of NLP7, promote its nuclear retention, or modulate its export dynamics.

Several other *nup* mutants are, like *nup98anup98b*, also modified for the intracellular partitioning of regulatory proteins, including TFs. For example, the mutation of the *HIGH EXPRESSION OF OSMOTICALLY RESPONSIVE GENES 1* (*HOS1*) reduces the nuclear accumulation of SnRK1a1 (Margalha et al., 2023) and of PHYTOCHROME-INTERACTING FACTOR 4 (PIF4) (Zhang et al., 2020). The NUP62-subcomplex modulates Flowering locus C (FLC) nuclear import, and NUP85 is a player in the nuclear-cytosolic trafficking of ACQOS, encoding a nucleotide-binding leucine-rich repeat (NLR) protein (Mori et al., 2024). The nuclear-cytosolic partitioning of these proteins is regulated by abiotic stresses or the transition to flowering. Here, we show that the response to nutrient availability is also governed by the function of a NUP, NUP98A/B.

*nup98anup98b* double mutants display early flowering (Jiang et al., 2019) with early senescence during flowering. NUP98A/B are also involved in the control of starch degradation (Xiao et al., 2020) and in the circadian control (Jiang et al. 2019, 2020). Here, we found that single *nup98a* mutants had only minor phenotypes, whereas the *nup98anup98b* double mutants showed a severe growth phenotype associated with early senescence as reported before (Xiao et al., 2020). Interestingly, the early senescence is exacerbated by N limitation. *nlp7* mutants display several traits of N limitation, despite growing under non-limiting nitrate supply (Castaings et al., 2009). Whereas this does not lead to early senescence of the *nlp7* mutants, impaired nuclear accumulation of other NLPs in the *nup98anup98b* mutant might lead to its early senescent phenotype under N limitation. Indeed, our study shows that in *nup98a-1nlp7-1* double mutants, a subset of nitrate-regulated genes displayed reduced transcript abundance in response to nitrate resupply compared to the nlp7 mutant. This might be due to the impact of the loss of function of NUP98A on the activities of other TFs involved in nitrate signaling, including NLP1/2/6/7 (Konishi et al., 2021, Cheng et al., 2023).

Besides the pivotal role in gating nuclear and cytoplasmic functions, the nuclear pore proteins function in key biological processes, including chromatin tethering to the nuclear periphery, a mechanism contributing to the regulation of gene expression (Ungricht and Kutay, 2017; Smith et al., 2015). Thus, it will require further investigations to determine whether NUP98A/B act with NLP to orchestrate the nitrate-triggered transcriptional regulation of nitrate-regulated genes by facilitating their transcriptional activation by NLPs in addition to regulating nuclear accumulation of NLPs in response to nitrate.

Interestingly, in humans, NUP98 has oncogenic roles as a genetic fusion partner (Gough et al., 2011). Indeed, in malignant cells, NUP98 occurs as part of fusion proteins with TFs, pointing to the involvement of NUP98 in transcriptomic dysregulation during oncogenesis. In plants, such NUP98 fusion proteins have not been reported, and it can be speculated that instead, protein-protein interactions evolved. Further studies are required to reveal further regulatory mechanisms governed by the NUP98-NLP7 protein complex.

In conclusion, this study demonstrates that in Arabidopsis, NUP98A/B promotes NLP7 activity through the control of its nucleocytoplasmic trafficking and thereby impacts the rapid transcriptional response to nitrate availability and the acclimation of plant growth to nitrate supply.

## Materials and Methods

### Plant materials

All Arabidopsis material is in the Col-0 background. The *Arabidopsis thaliana* mutant *nlp7-1* (SALK_26134) derived from T-DNA–mutagenized populations of the Col-0 accession (Alonso-Blanco et al., 2003, Castaings et al., 2009) and was backcrossed twice to Col-0. All mutant seeds were obtained from the NASC, and homozygous lines were identified by genotyping using the primers listed in Supplemental Table S1. Homozygote lines of *nup98a1* (SALK_103803, Jiang et al., 2019), *nup98a2* (SALK_015016, Jiang et al., 2019, also named *dra2-4*, Gallemi et al., 2016), *nup98b1* (GABI_288A08, Jiang et al, 2019), and *nup98b2* (SALK_132527) have been selected by genotyping. The higher-order mutants with and without the *NLP7pro:GFP-NLP7* construct have been obtained by crosses of *NLP7pro:GFP-NLP7* (Col-0) line A4.4 (Castaings et al., 2009) followed by selfing of the F1 generation, genotyping the F2 generation and selecting for homozygous transgene by hygromycin selection in F3.

### Bimolecular Fluorescence Complementation (BiFC)

Open reading frames (without stop codon) flanked by AttB1 and AttB2 sites were amplified from Arabidopsis cDNA clones (Columbia accession) using the primers as given in Supplemental Table S1, cloned into Gateway vector pDONR207 using BP recombination (Invitrogen), and sequenced, except *NUP98A*, where the cDNA sequence optimized for *Arabidopsis thaliana* was synthetized and cloned into pTwist ENTR Kozak entry vector by Twist Biosciences (Supplemental Data Set 1). An LR reaction between the entry vector and the complete set of four pBiFP vectors (Azimzadeh et al., 2008) produced the final expression vectors, where coding sequences are cloned in fusion with the N- or C-terminal parts of YFP, either as N-terminal or C-terminal fusions, under the control of the cauliflower mosaic virus 35S promoter.

Modified versions of NLP7 were produced by PCR. For NLP7 without PB1 domain, the region corresponding to 1-863 amino acids was amplified by PCR and cloned into the Gateway vector pDONR207 using BP recombination. For NLP7 without the predicted GAF domain (Chardin et al., 2014), regions corresponding to amino acids 1-159 and 301-959 were amplified separately. The two fragments were fused together by a second PCR, and the resulting fragment flanked by AttB1 and AttB2 sites was cloned into pDONR207. The primer sequences are given in Table S1. A description of the pH2B2:H2B2-mTurquoise construct can be found in Bouchez et al. (2024).

For BiFC interaction assays, the vectors were transformed into *Agrobacterium tumefaciens* strain C58C1(pMP90) (Kámán-Tóth et al., 2018) and co-infiltrated into *Nicotiana benthamiana* leaves. Plants were grown either in fertilized soil in an air-conditioned greenhouse or on sand in a growth chamber under long day conditions (day 22 oC / night 18 °C, 16h photoperiod, 130-160 µmolm-2s-1 light intensity (Systion 32-42 VDC 330mA, 11V). Plants grown in sand culture were watered with a 2 mM nitrate-containing nutrient solution for 29 days, followed by watering for 3 days with a nutrient solution without N after washing the sand with the N-depleted solution or by watering with a 10 mM nitrate nutrient solution. Agrobacterium carrying clones of interest were grown overnight at 28°C in 5 ml LB medium with appropriate antibiotics. Aliquots from the overnight cultures were resuspended in 10 mM MgCl_2_ and 10 mM 2-(N-morpholine)-ethanesulphonic acid (MES), pH 5.6, to obtain a final OD600nm of 0.5 for tobacco leaf infiltration. YFP fluorescence was detected 2-3 days after infiltration. During that time, plants were watered with either 0 mM or 10 mM nitrate-containing nutrient solution. Ion equilibrium of the nutrient solution was ensured by replacing KNO3 and Ca(NO_3_)_2_ with KCl and CaSO_4_, respectively.

### Yeast Two-Hybrid Assays

Coding sequences of NLP7 and NUP98A cloned in pDONR207 were recombined into the pGBKT7 and pGADT7 vectors (Clontech) via LR Clonase reactions and co-transformed into the Saccharomyces cerevisiae strain MaV203 (Invitrogen). Yeast was cultured on YPDA plates (Clontech) at 28 °C. For transformation, an overnight liquid culture at OD600nm =0.6 was harvested, resuspended in 1 mL sterile water, pelleted by centrifugation (maximum speed, 15 s), and resuspended in 1 mL of 0.1 M lithium acetate in 1X TE buffer. Transformation mixes contained 5 µg of each plasmid, 5 µL of carrier DNA (Finnzymes, Fisher Scientific), 0.1 mL of yeast suspension, and 0.6 mL of 40% [v/v] PEG 4000 in 0.1 M lithium acetate/1X TE. After a 30 min incubation at 28 °C, mixtures were heat-shocked at 42 °C for 25 min. Cells were pelleted (15 s, max speed), resuspended in 0.25 mL sterile water, and plated on SD/-Trp-Leu selective medium (Clontech). To assess protein–protein interactions, cotransformed colonies were resuspended in 0.1 mL sterile water, and 10 µL was spotted on SD/-Trp-Leu-His plates supplemented with various concentrations of 3-amino-1,2,4-triazole (3-AT). Growth under these conditions was interpreted as a positive interaction.

### Growth conditions of Arabidopsis plants

Seeds were stratified for 5 days at 4°C in the dark and then grown either on non-fertilized soil (Stender) or on sand, both supplemented with nutrient solutions (Loudet et al., 2003) containing either 0.5 mM (growth-limiting nitrate supply) or 5 mM (non-limiting nitrate supply) nitrate, in short-day photoperiod with an 8-h light/16-h dark cycle at 21°C/18°C, respectively, 65% relative humidity, and a constant 150 μmol photons m^−2^ s^−1^ irradiation with Osram 36w light sources. For N limitation conditions, ion equilibrium of the medium was ensured by partly replacing KNO_3_ and Ca(NO_3_)_2_ with KCl and CaSO_4_, respectively.

For nitrate response assays, both for the determination of steady state mRNA levels and confocal analyses, seeds were stratified for 3 days at 4°C and then grown in liquid medium containing 0.07% (w/v) MES pH 5.8, 1 mM KH_2_PO_4_, 0.5 mM MgCl_2_, 0.5 mM CaSO_4_, microelements as in Estelle and Somerville (1987), 0.25% (w/v) sucrose and 3 mM KNO_3_ for 10 days in long-day photoperiod (16-h light/8-h dark) at 25°C with a light intensity of 80 µmol photons ^.^ m^−2.^s^−1^ by Philips F17T8/TL741 light sources. For gene expression analysis after 7 days of growth, the medium was changed to fresh medium every 2 days. Eleven-day-old seedlings were then nitrogen-starved for 3 days using fresh medium containing 3 mM KCl instead of 3 mM KNO_3_. After 3 days of N starvation, 3 mM nitrate was added for the indicated time periods, and harvests were performed at several time points. For confocal analysis, seedlings were grown as in Marchive et al. (2013) for five days on a 3 mM KNO_3_ liquid medium, then N-starved for two days by exchanging the culture medium with an N-free medium in which KNO_3_ was replaced by KCl.

### Confocal Imaging

Laser scanning confocal imaging was performed using the SP8 confocal laser microscope (Leica, Germany) equipped with an argon laser (488 nm for GFP excitation, intensity 20%) and an argon laser (514 nm for YFP excitation, intensity 20%). Emission was collected at 505 to 525 nm (GFP, gain= 500) or at 520 to 540 nm (YFP, gain= 500). The images were coded green for both GFP and YFP. Images were processed with ImageJ. Plant samples were mounted in 40 µL of a top agar solution (0.1%, w/v). During confocal imaging, 5 µL of 0.5 M KNO_3_ was added by capillarity to the mounting medium (45 µl).

### Gene expression analysis

Measurements of relative steady-state mRNA levels were performed on seedlings (plants at 1.02 developmental growth stages (Boyes et al., 2001)) that had been cultivated in liquid medium as described above. Each sample was shock frozen after harvest and was composed of 26-34 seedlings. Total RNA was extracted using the RNeasy plant mini kit (QIAGEN), including a DNase treatment according to the manufacturer’s instructions, or by Trizol method (Invitrogen). First-strand cDNAs were synthesized according to Daniel-Vedele and Caboche (1993) using Moloney murine leukemia virus reverse transcriptase and oligo(dT)15 primers (Promega) or using the SuperScript III Fast strand synthesis system (Invitrogen).

cDNAs were first checked by PCR for contamination by genomic DNA, then relative expression was determined in 384-well plates with a QuantStudio 5 System device (40 3-step cycles; step 1, 95°C, 5 s; step 2, 58°C, 30 s; step 3, 72°C, 30 s) using Takyon qPCR kit for SYBR assay (Eurogentec) (2.5 µL TAKYON SYBR 2X, 0.015 µL Forward primer 100 µM, 0.015 µL Reverse primer 100 µM, 2.5 µL cDNA 1/20 per well). Relative expression was calculated according to the 2-ΔCt method. Target gene expression was normalized to the geometric mean of the expression of two plant genes, *ACTIN2* (At3g18780) and *TIP41* (At4g34270). All primer sequences are given in Supplemental Table S1.

### Statistical analyses

Data were analyzed using non-parametric tests due to non-normal distributions and/or unequal variances. Comparisons between multiple groups were performed using the Kruskal–Wallis test. When a significant difference was detected (p < 0.05), Dunn’s post hoc test was used for pairwise comparisons without correction for multiple testing. Statistical analyses were conducted in Python using the scipy and scikit-posthocs libraries.

## Acknowledgments

We thank Hervé Ferry, Jean-Christophe Guedet and Philippe Maréchal for plant care, Gladys Cloarec for support for confocal analyses, and Alizée Teinturier for technical assistance. We also thank the NASC and ABRC for supplying Arabidopsis T-DNA insertion lines. This work has benefited from the support of IJPB’s Plant Observatory technological platforms PO-Plants and PO-Cyto and was supported by ANR NITRASENSE (ANR-16-CE20-022, ZK, AK). The IJPB benefits from the support of the Saclay Plant Sciences (SPS) Network (ANR-17-EUR-0007). We thank Sylvie Ferrario-Méry, Thomas Girin, Anne-Sophie Leprince and Christian Meyer for fruitful discussions.

## Author Contributions

Z.K and A.K. designed research; Z.K, G.D., V.B., and A.K. performed research and analyzed data; and Z.K., G.D., and A.K. wrote the paper.

---

**Supplemental Table S1.**
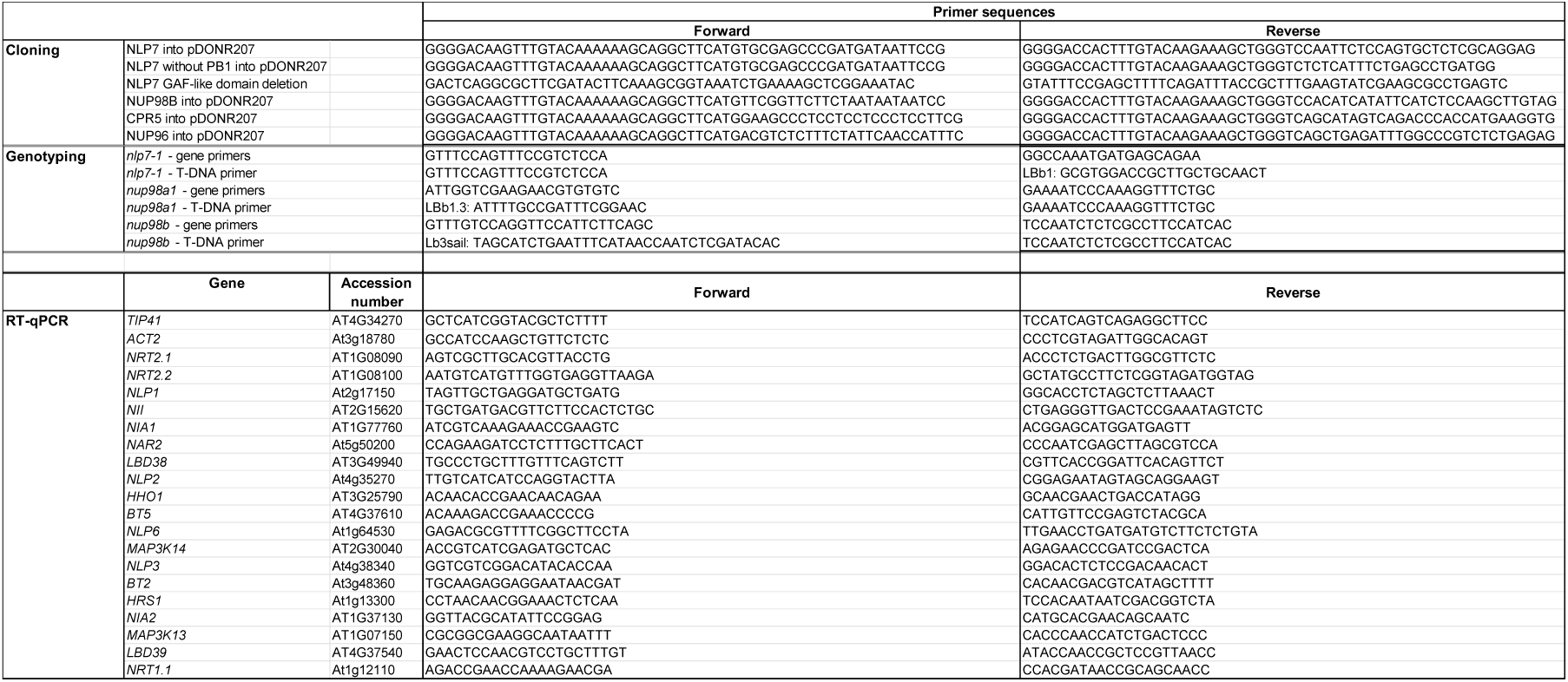
Primers used for cloning, genotyping and RT-qPCR analyses.

**Supplemental Figure S1.**
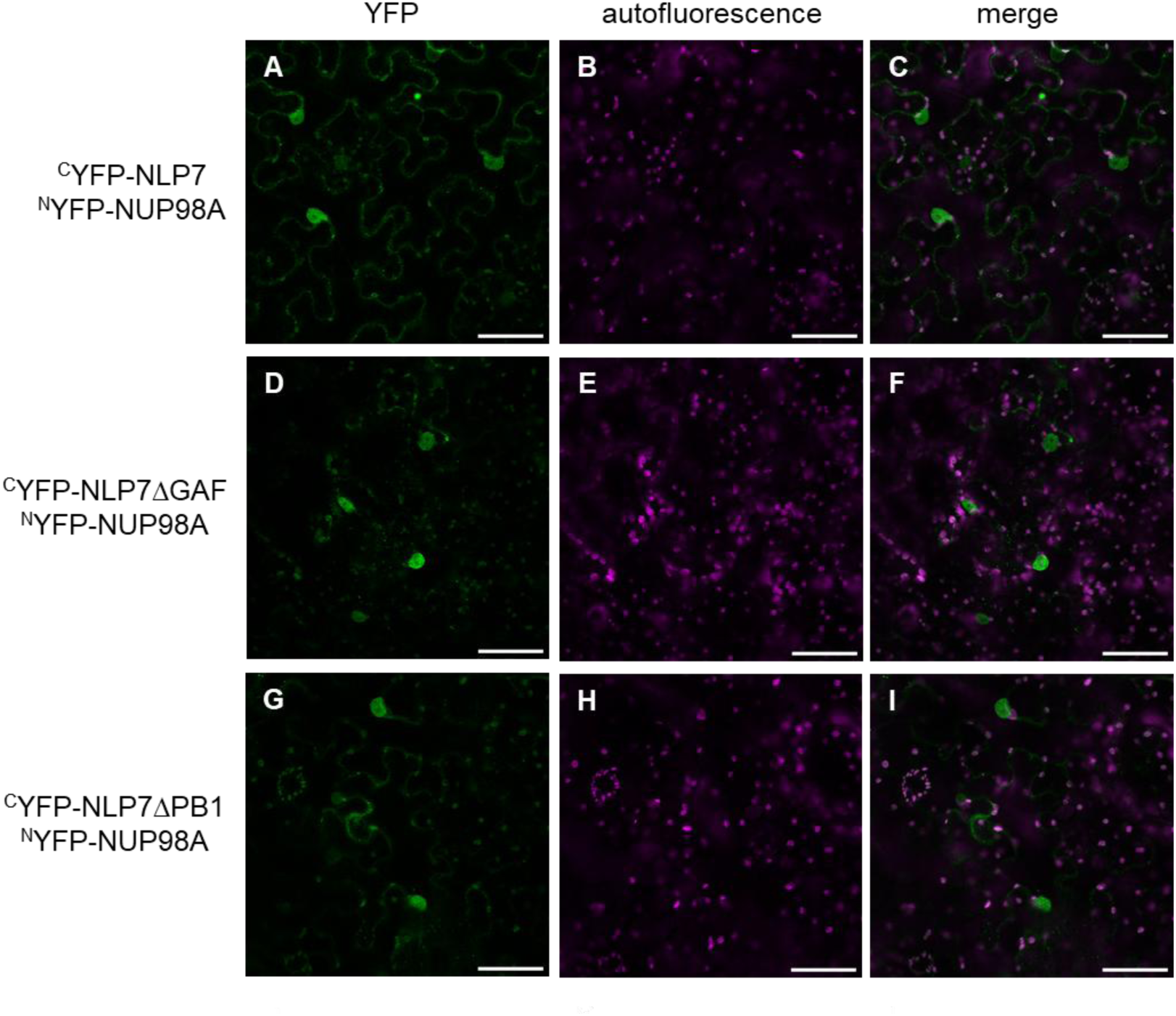
NLP7 GAF and PB1 domains are not essential for the NLP7-NUP98A interaction. NLP7ΔGAF and NLP7ΔPB1, interact with NUP98A in BiFC assays in *N. benthamiana* (**A-C**) NLP7-NUP98A interaction, (**D-F**) NLP7DGAF-NUP98A interaction, **(G-I)** NLP7DPB1-NUP98A interaction. (**A, D, G**) YFP fluorescence (green), (**B, E, H**) chlorophyll autofluorescence (magenta) (**C, F, I**) YFP fluorescence (green) was merged with chlorophyll autofluorescence (magenta). Scale bar = 50 µm.

